# Arginine culture generates triacylglycerol by triggering nitrogen starvation responses during robust growth in *Chlamydomonas*

**DOI:** 10.1101/416594

**Authors:** Jacob Munz, Yuan Xiong, Jaoon Young Hwan Kim, Young Joon Sung, Seungbeom Seo, Ran Ha Hong, Thamali Kariyawasam, Nolan Shelley, Jenny Lee, Sang Jun Sim, EonSeon Jin, Jae-Hyeok Lee

## Abstract

Under nitrogen (N) starvation, microalgae increase carbon storage in the form of lipid droplets while also downregulating photosynthesis and eventually terminating growth. To improve lipid yield, we asked whether lipid droplets and N starvation responses can be induced without limiting growth or photosynthesis. In the chlorophyte *Chlamydomonas reinhardtii*, gametogenesis is induced either by N starvation or by growth with arginine as the sole N source. We showed that arginine cultures supported robust phototrophic growth, constitutively turned on N starvation-induced genes, and increased lipid droplets. The lipids accumulated in arginine cultures exhibited strong enrichment of saturated and monounsaturated fatty acids, a preferred characteristic of biodiesel precursors. The diatom *Phaeodactylum tricornutum* also accumulated lipid droplets in arginine culture without growth impairment. We document a system wherein N starvation responses are induced without compromising photosynthesis or growth, thereby suited to the producing valuable chemicals and biofuel precursors without requiring stressors in microalgae.

## 1. Introduction

Microalgae are a promising feedstock for biofuels and other valuable biocompounds compared to traditional crop systems due to their potentially superior productivity, oil content, carbon dioxide sequestration, and ability to be cultured on non-arable farmland (Antizar-Ladislao and Turrion-Gomez, 2008). Additionally, microalgal triacylglycerides have a short harvesting cycle -- less than one month -- compared with vegetable oils from food crops (Schenk et al., 2008). Indeed, microalgae could potentially produce up to 17 times more oil per hectare of culture than even the most productive vegetable oil crops (Robles-Medina et al., 2009).

Microalgae represent a suitable replacement for vegetable crop biodiesel precursors only if their oil contains large amounts of fatty acid (>40% w/w) (Chisti, 2007). Biodiesel derived from microalgae low in polyunsaturated fatty acids possesses a higher cetane number than biodiesel obtained from other sources that are rich in polyunsaturated fatty acids (James et al., 2013). Therefore, enrichment in saturated and monounsaturated fatty acids such as C12:0, C14:0, C16:0, C16:1, C18:0, and C18:1 (carbon number : number of double bonds) is desirable for microalgal feedstocks (Chisti, 2007).

The lipids derived from microalgae are predominantly stored in lipid droplets rich in triacylglycerols (TAGs) whose biosynthesis is complex. The model unicellular algae *Chlamydomonas reinhardtii* is estimated by sequence analysis to possess 113 genes related to lipid metabolism whereas the model diatom *Phaeodactylum tricornutum* is estimated to have 106 genes (Li-Beisson et al., 2015; Mühlroth et al., 2013). The usual protocol to stimulate lipid accumulation in microalgae is by way of nutrient limitation, most commonly nitrogen (N) (Hu et al., 2008; Wang et al., 2009). However, N starvation-induced lipid production is accompanied by a strong decrease in overall biomass since N is essential for growth, impeding the goal of economical microalgal lipid cultivation.

N limitation responses are triggered by signaling mechanisms that sense external and internal N availability. Transcriptional/proteomic changes stimulated by N starvation in *C. reinhardtii* have been intensively studied (Schmollinger et al., 2014; Wase et al., 2014). N limitation rapidly induces genes including the common N catabolite genes for N scavenging and N salvaging, suggesting that *C. reinhardtii* actively remobilizes N from purines and amino acids while searching for alternative external N sources (Park et al., 2015; Schmollinger et al., 2014). N limitation also downregulates photosynthesis genes including the majority that encode antenna and photosystem proteins (39 out of 42 genes in Miller et al., 2010).

Physiological responses to N limitation in *C. reinhardtii* include the transition to sexually competent gametes (gametogenesis), degradation of photosynthetic proteins/chlorophylls, ribosome turnover, and the accumulation of carbon storage molecules (Bulté and Wollman, 1992; Martin and Goodenough, 1975; Sager, 1954; Siersma and Chiang, 1971). Gametogenesis is a response unique to N starvation: the removal of any single component other than N from the growth medium does not elicit gametogenesis, and cultures produce <1% sexually competent gametes when the preferred N sources, ammonium or nitrate, are present (Matsuda et al., 1992; Pozuelo et al., 2000; Sager and Granick, 1953).

*C. reinhardtii* can grow on various N sources ranging from simple ammonium and amino acids to complex nucleic acids and their derivatives (Sager and Granick, 1953). Muñoz-Blanco et al., (1990) reported that twelve amino acids can serve as the sole N source for *C. reinhardtii* growth, but in all but one case, these amino acids undergo deamination via an extracellular L-amino acid oxidase encoded by *LAO1*, releasing ammonium which then enters the cell. Arginine, the exception, is reported to enter the cells by a specific active transporter system (Kirk and Kirk, 1978) whose molecular identity has been proposed but not determined experimentally (Vallon and Spalding, 2009). Interestingly, arginine is the only N source reported to allow gamete production while retaining a growth rate comparable to ammonium or nitrate (Honeycutt and Margulies, 1972; Matsuda et al., 1992). These studies suggest that the occurrence of gametogenesis in arginine-grown populations is not due to a lack of N but rather to the lack of a repressive gametogenesis signal such as ammonium.

With the goal of improving the lipid yields in microalgae by manipulating N starvation responses, we report in this study the physiological and molecular responses of *C. reinhardtii* cultures grown in arginine-based media. We document that arginine cultures exhibit multiple N starvation-induced responses including the accumulation of lipid droplets without compromising growth and photosynthesis.

## 2. Materials and methods

### 2.1. Strains and culture conditions

*Chlamydomonas reinhardtii* wild-type strains 137c (CC-125: *nit1; nit2; mt*+) and 21gr (CC-1690: *NIT1; NIT2; mt*+), field-isolated strains CC-2344 (J356, *mt*+), CC-2290 (*mt*-), CC-2932 (*mt*+), and CC-2931 (*mt*-), a starch-less mutant *sta6* (CC-4348), and mating tester strains CC-620 (*mt*+) and CC-621 (*mt*-) were obtained from the Chlamydomonas Resource Center (https://www.chlamycollection.org/). *NIT1* and *NIT2* genotypes of 137c and 21gr were confirmed by nitrate- and nitrite-dependent growth phenotypes and sequencing of *nit2* alleles. All strains were maintained under low light (30 μmol photons m^−2^ s^−1^) at 23°C in Tris-Acetate-Phosphate (TAP) medium, unless otherwise stated. N-free TAP medium was prepared by omitting N from TAP medium. Experimental cultures were grown under 100 μmol photons m^−2^ s^−1^ at 23°C on an orbital shaker. Phototrophic cultures were grown under 150 μmol photons m^−2^ s^−1^ at 23°C in TAP medium omitting acetate.

Considering the four-fold difference in N content between ammonium and arginine, we tested various concentrations of ammonium vs. arginine in TAP media and determined that 2 mM arginine and 4 mM ammonium was sufficient for maximum growth. To ensure sufficient N supply, we used 8 mM arginine or 8 mM ammonium (hereafter referred to as arginine cultures or ammonium cultures), concentrations that produced comparable growth rates and yield to the established minimum concentrations.

Growth curves were generated using a small scale photobioreactor (Multi-Cultivator MC1000; Photon System Instruments, Czech Republic) measuring the absorption at 680 nm every five minutes with an initial cell concentration of 2 x 10^5^ cells per mL. Cultures were grown under 100 μmol photons m^−2^ s^−1^ at 23°C and bubbled with ambient air (mixotrophic) or grown under 250 μmol photons m^−2^ s^−1^ at 23°C and bubbled with 5% CO2 mixed with air (phototrophic). Growth curve measurements using optical density can be influenced by the chlorophyll contents of the culture; therefore, cell number was monitored manually using a hemocytometer following fixation of cells in 2% glutaraldehyde. Growth rates and doubling times were calculated as in Widdel (2007).

An axenic stock of the diatom *Phaeodactylum tricornutum* Bohlin (CCMP632) was obtained from the National Center for Marine Algae and Microbiota at the Bigelow Laboratory for Ocean Sciences. Cells were cultured in f/2+Si medium (enriched artificial seawater) containing 0.88 mM sodium nitrate and 10 mM sodium bicarbonate. For arginine culture, sodium nitrate was replaced by the same N content of arginine (0.22 mM). Cultures were grown under fluorescent lamps (50 μmol photons m^−2^ s^−1^) on an orbital shaker at 20°C with a 12h/12h light/dark regime.

### 2.2. Measurement of gametogenesis

Ten-day-old plate-grown cells were resuspended in 1 mL of liquid ammonium or arginine TAP to a final density of 2 x 10^6^ cells per mL and incubated in 24-well plates for 24 hours before mating efficiency analysis. Mating efficiency was measured with the CC-620 or CC-621 tester gametes that were incubated for six hours in N-free TAP medium as described in Galloway and Goodenough (1985).

### 2.3. *Reverse transcription quantitative PCR (RT-qPCR)* of *N starvation marker genes*

The eight N starvation marker genes used in this study are involved in N scavenging (*DUR3A, NRT2.1, NIT1*, and *LAO1*) and N salvaging (*AMT4, AIH2, FDX2* and *GLN3*), and were previously shown to be upregulated between 10- and 1000-fold within two hours of N starvation with stably low basal expression in N replete condition (Schmollinger et al., 2014). Details of the marker genes are summarized in Table 1.

**Table 1.**
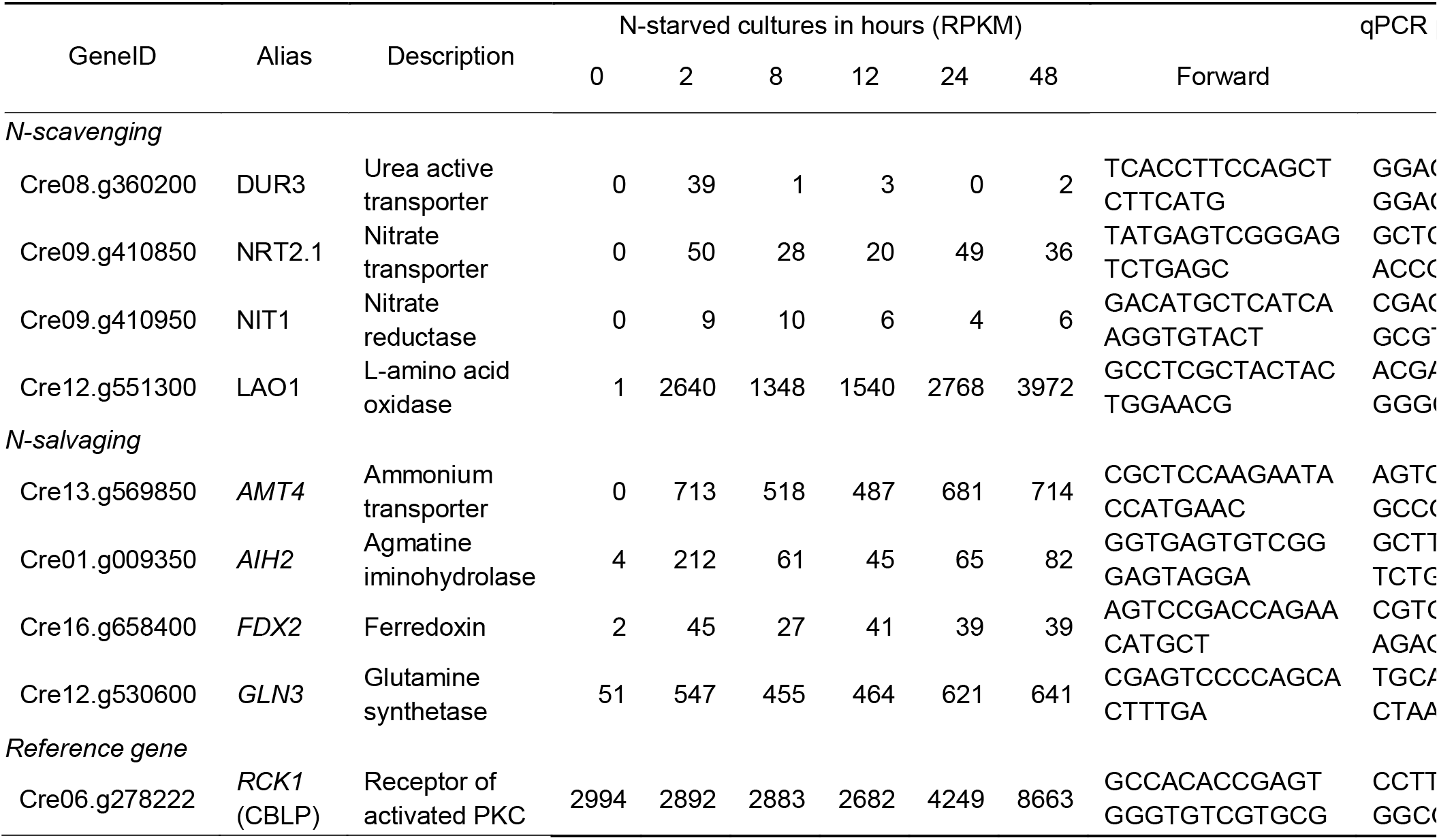
Details of selected N-starvation marker genes analyzed in this study. Transcriptome data of mixotrophic cultures is from Schmollinger *et al.* (2014), which is also available on www.phytozome.net

For analyzing transcript accumulation, mid-logarithmic phase cultures were harvested at 3,000 rpm for 3 minutes and washed twice with 50 mL N-free medium. Washed cells were added to 25 mL of arginine, ammonium, or N-free medium to a final concentration of 2 x 10^6^ cells per mL and incubated for 2 hours or 24 hours. RNA extraction and RT-qPCR were performed as previously described (Joo et al., 2017). Results were obtained in technical duplicates for three biological replicates and quantified by normalization to the endogenous reference gene *RCK1* (also known as *CBLP*) that has been commonly used in *C. reinhardtii* (Allen et al., 2007). Primer sequences and amplicon lengths are presented in Table 1.

### 2.4. Chlorophyll a fluorescence and pigment determination

*In vivo* chlorophyll fluorescence was measured with a JTS-10 spectrophotometer of 100 μL of cells at a concentration of 5 x 10^6^ cells per mL in a stainless steel plate (Bio-Logic Science Instruments, France) and recorded with a CCD camera (C9300-221, Hamamatsu, Japan). Samples were incubated in the dark for 10 minutes prior to fluorescence measurements. Maximum quantum efficiency of PSII photochemistry (F_v_ / F_m_) and PSII operating efficiency (Φ_PSII_) under an actinic light intensity of 210 μmol photons m^−2^ s^−1^ were assayed and calculated according to Baker (2008).

Chlorophyll was extracted in 80% acetone and cell debris removed by centrifugation at 20,000 *g* for 5 minutes. Chlorophyll was quantified by examination of supernatant light absorbance in a cuvette at 645 nm, 665 nm and 720 nm with a spectrophotometer (DU 730, Beckman Coulter, USA) (Arnon, 1949) with equations corrected as in Melis et al. (1987).

### 2.5. Visualization of lipid droplets

Lipid droplets were stained with addition of Nile red stock (0.1 mg per mL in acetone) (Sigma-Aldrich, USA) at a final concentration of 1 μg per mL followed by a 20 min incubation in the dark at room temperature. A 488 nm excitation and a 560 to 600 nm emission wavelength were used to capture the Nile red signal. Images of *C. reinhardtii* cells were captured with a Leica DFC350 FX camera mounted on a Leica DM6000 B microscope with a 100x oil immersion lens objective. *P. tricornutum* lipid droplets were also stained by the same method described above. 1% low-melting agarose was mixed with the cultures (1:1); and stained cells were observed under a laser-scanning confocal microscope, ECLIPSE Ti (Nikon Corp.) using a 561 nm excitation wavelength and a long-pass 570 nm emission filter. The images depict representative cells.

### 2.6. Analysis of neutral lipid content

Total lipids were extracted according to Bligh and Dyer (1959), and as described in more detail (Kim et al., 2016). Cell suspensions were concentrated to 1 x 10^8^ cells per mL. Cellular and subcellular fractions were extracted with methanol-chloroform-formic acid mixture (2:1:0.1, v/v) and phases were separated by adding 0.5 mL of 1 M potassium chloride and 0.2 M phosphate. The organic phase was loaded on silica gel matrix aluminum plates (20 x 20 cm, Sigma-Aldrich, USA) immersed in 0.15 M ammonium sulfate for 30 minutes and dried for at least 2 days. Lipids were separated on thin layer chromatography plates using a double development solvent system (2/3 in acetone-toluene-water (91:30:3, v/v)) and then fully in hexane-diethyl ether-acetic acid (70:30:1, v/v) in a sealed container. TAGs were identified using the soybean lipid profile as a reference and scraped off the plates. The scraped lipids and silica were transferred to a screw-cap glass tube and 1 mL of hexane was added. For gas chromatography, all samples were methylated by acid-catalyzed transesterification using methanol with sulfuric acid (3%, v/v) and heated at 95°C for 1.5 hours (Lim et al., 2014), and pentadecanoic acid (C15:0) was added to 1 mg per mL as an internal standard. After collection of the organic phase, FAME was analyzed using gas chromatography (Agilent 7890A) with a flame ionization detector and a DB-23 column (Agilent, USA) with the following protocol: injection volume, 1 μL; split ratio, 1:50; inlet temp, 250°C; detector temp, 280°C; oven temp, hold at 50°C for 1 min, increase to 175°C at 25°C min^−1^, increase to 230°C at 4°C min^−1^, and hold for 5 min. TAGs and FAMEs were quantified based on the number of cells extracted.

*P. tricornutum* lipid quantitation with Nile red was performed as described with modifications (Chen et al., 2009). Cells were permeabilized with 20% DMSO for 20 min followed by the addition of 10 μL Nile red stock (0.1 mg per mL in acetone) to 1 mL of culture, inverting rapidly. Samples were incubated in the dark for 20 min at room temperature. 200 μL of stained culture was then transferred to a black well plate and fluorescence was detected using a Varioskan Flash spectral scanning multimode reader (Thermo Fisher Scientific, USA) with a 530 nm excitation and a 580 nm emission wavelength. The measurement was divided by cell concentration, determined using a hemocytometer, to ascertain relative fluorescence levels.

## 3. Results and Discussion

### 3.1. N starvation-induced genes are upregulated in arginine cultures that undergo gametogenesis and reduce chlorophyll content during normal growth

Previous reports documenting gametogenesis in arginine cultures characterized the 137c strain which is defective in nitrate assimilation (Honeycutt and Margulies, 1972; Matsuda et al., 1992). To test whether the gametogenesis induced in arginine cultures depends on the 137c-specific genetic background, we characterized two strains, 21gr (CC-1690: *NIT1; NIT2*) and 137c (CC-125: *nit1; nit2*), where *NIT1* encodes nitrate reductase (Fernández and Matagne, 1984) and *NIT2* encodes the transcription factor regulating nitrate assimilation genes (Fernández and Matagne, 1986). In 21gr and 137c about ~10% of the population were gametes in arginine cultures in contrast to little or no gametes in ammonium cultures (Table 2a). We extended the query to four natural isolates of *C. reinhardtii* whose genomes contain 2 - 3% difference at the nucleotide level with the laboratory strains such as 21gr and 137c (Flowers et al., 2015). Arginine cultures of the natural strains exhibited 5 - 35% of gametes (Table 2a), indicating that arginine culture induces gametogenesis in diverse genetic backgrounds of *C. reinhardtii.*

**Table 2.**
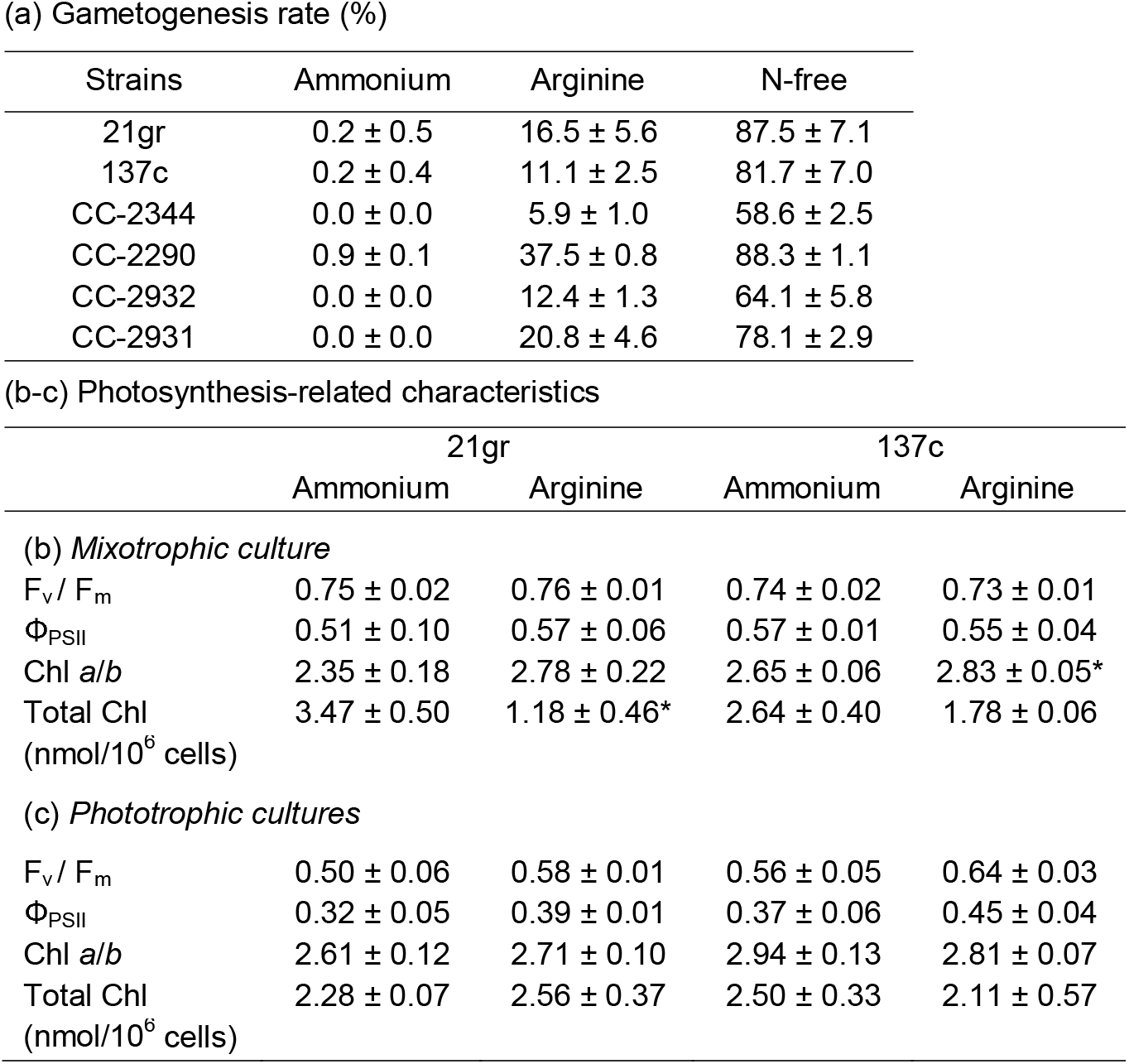
*C. reinhardtii* cells acclimated to arginine media exhibit N-starvation-induced responses without stress symptoms. (a) Mating efficiency was used to measure gametogenesis rate, determined by the percentage of cells fused with tester gametes. Data represents the mean ± the standard deviation from three biological replicates, except for field-isolated strains examined by single experiment with two replicates. (b-c) Properties of *C. reinhardtii* cells acclimated to different nitrogen sources. Cells from mid-logarithmic mixotrophic (b) or phototrophic (c) cultures used to determine the F_v_ / F_m_, Φ_PSII_ at 210 μmole photons m^−2^ sec^−1^, chlorophyll *a/b* ratio, and total chlorophyll. Values shown are means ± standard deviation (n=2). Significance is indicated with an asterisk and was calculated by a Student’s t-test (P < 0.05). Chl, chlorophyll.

Using a photobioreactor we examined growth curves of arginine and ammonium cultures, measured as optical density at 680 nm (Fig. 1a). Doubling time calculated from the growth curves was 14 – 21% increased in the arginine cultures compared to the ammonium cultures during the logarithmic phase (Table 3a). Manual cell counting showed that arginine cultures had a significantly higher maximal cell density at 1.97 ± 0.19 x 10^7^ cells per mL in 21gr and a comparable maximal cell density at 1.35 ± 0.13 x 10^7^ cells per mL in 137c to ammonium cultures (Table 3a). Overall, arginine cultures showed 12.5 - 16.5% slower growth rates but contained comparable or 1.5 times more cells per volume when they reached stationary phase (Table 3a).

**Fig. 1.**
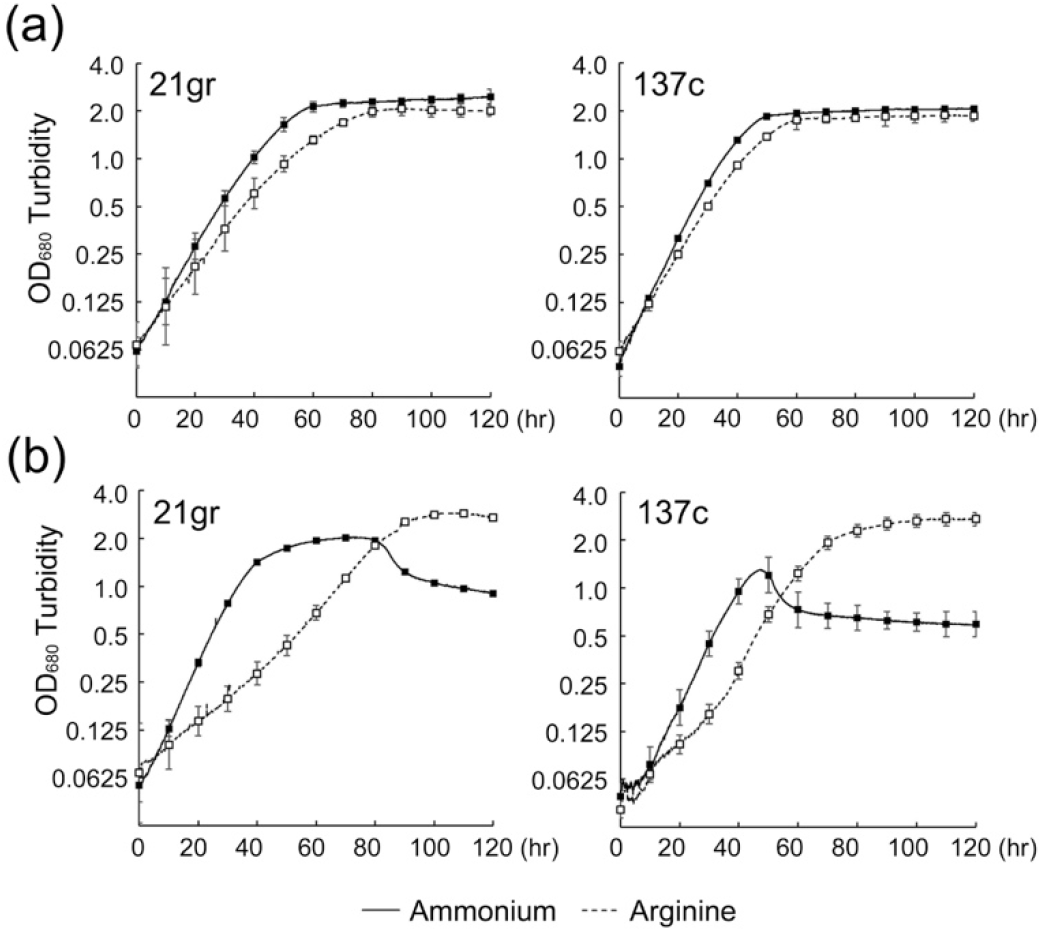
Arginine cultures grow slower than ammonium cultures but reach to a higher density when grown in phototrophic condition. Growth curves of 21gr and 137c cultures in mixotrophic condition (a) or in phototrophic condition (b), measured as optical density at 680 nm on a log_2_ scale. Solid and dashed lines indicate ammonium and arginine cultures, respectively. Error bars are standard deviation from two or more independent experiments.

**Table 3.**
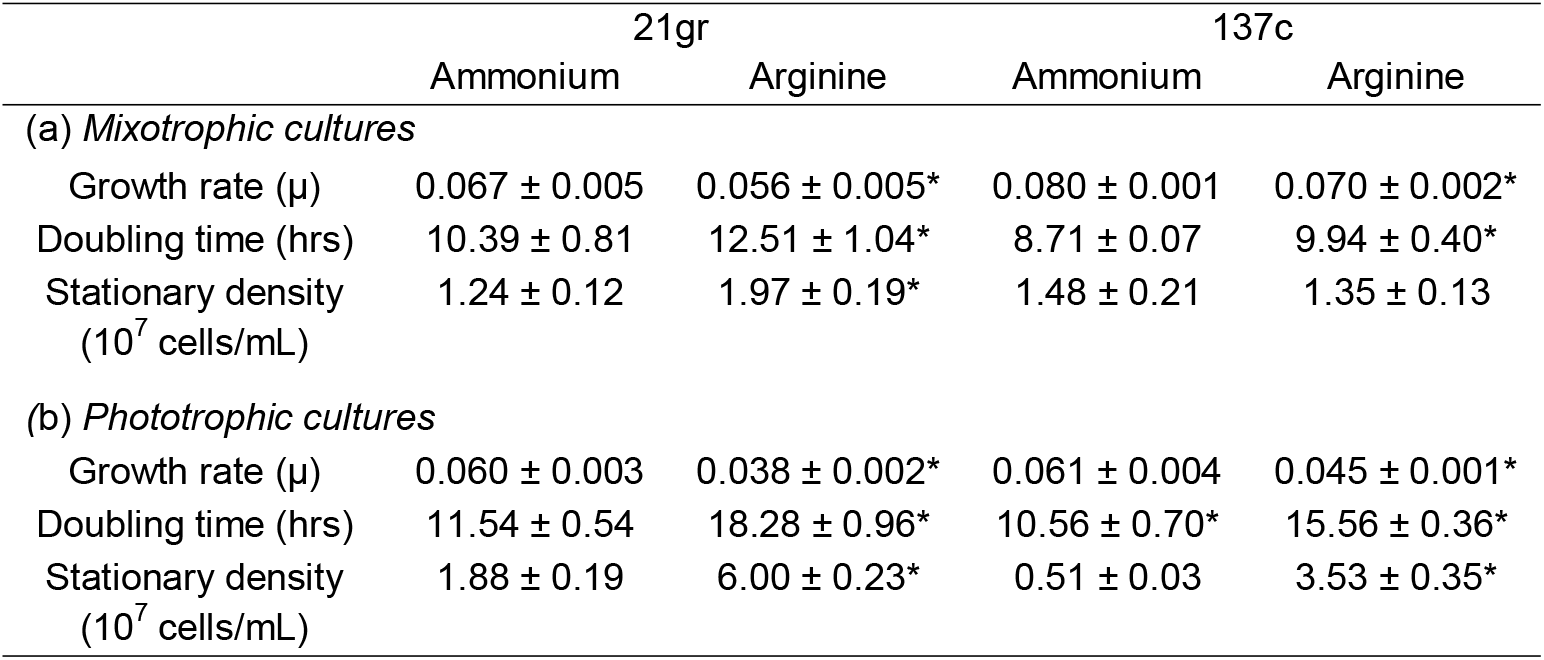
*C. reinhardtii* growth characteristics of ammonium and arginine cultures in photobioreactor. The growth rate (μ) and doubling times were calculated during ten hours of linear growth of the logarithmic phase in mixotrophic (a) and phototrophic (b) cultures. Values are means ± standard deviation (n=3). Significant difference from ammonium data is indicated with an asterisk and was calculated by a Student’s t-test (P < 0.05).

|  | 21gr |  | 137c |  |
| --- | --- | --- | --- | --- |
|  | Ammonium | Arginine | Ammonium | Arginine |
| *(a) Mixotrophic cultures* |  |  |  |  |
| Growth rate ($\mu$) | 0.067 ± 0.005 | 0.056 ± 0.005* | 0.080 ± 0.001 | 0.070 ± 0.002* |
| Doubling time (hrs) | 10.39 ± 0.81 | 12.51 ± 1.04* | 8.71 ± 0.07 | 9.94 ± 0.40* |
| Stationary density<br>(10<sup>7</sup> cells/mL) | 1.24 ± 0.12 | 1.97 ± 0.19* | 1.48 ± 0.21 | 1.35 ± 0.13 |
| *(b) Phototrophic cultures* |  |  |  |  |
| Growth rate ($\mu$) | 0.060 ± 0.003 | 0.038 ± 0.002* | 0.061 ± 0.004 | 0.045 ± 0.001* |
| Doubling time (hrs) | 11.54 ± 0.54 | 18.28 ± 0.96* | 10.56 ± 0.70* | 15.56 ± 0.36* |
| Stationary density<br>(10<sup>7</sup> cells/mL) | 1.88 ± 0.19 | 6.00 ± 0.23* | 0.51 ± 0.03 | 3.53 ± 0.35* |

One of the early responses to N starvation is the rapid upregulation of N catabolite genes within 0.5 - 2 hours after removal of preferred N sources (Schmollinger et al., 2014). To ask whether this response occurs in growing arginine cultures, we analyzed expression levels of eight N starvation marker genes involved in N scavenging and N salvaging (Table 1). All were upregulated between 10- and >1,000-fold in 2 hours after ammonium cultures were transferred to N-free medium (Fig. 2a, N-free of ammonium culture), and a comparable upregulation was observed when transferred to arginine medium (Fig. 2a, Arg of ammonium culture). The upregulated expression remained high for all marker genes in 21gr after 24 hours in arginine medium, whereas in 137c, *AIH2, DUR3A, LAO1* showed some reduction but were still 8-to 125-fold higher than the levels in ammonium culture, while *FDX2* and *NRT2.1* expression fell to the levels in ammonium culture (Fig. 2a, Arg-24hr of ammonium culture). The reduced 24-hour expression of *AIH2* and *DUR3A* in arginine medium is concordant with their transient upregulation within 2 hours and reduced expression in the next hours of N starvation (Schmollinger et al., 2014; Table 1).

**Fig. 2.**
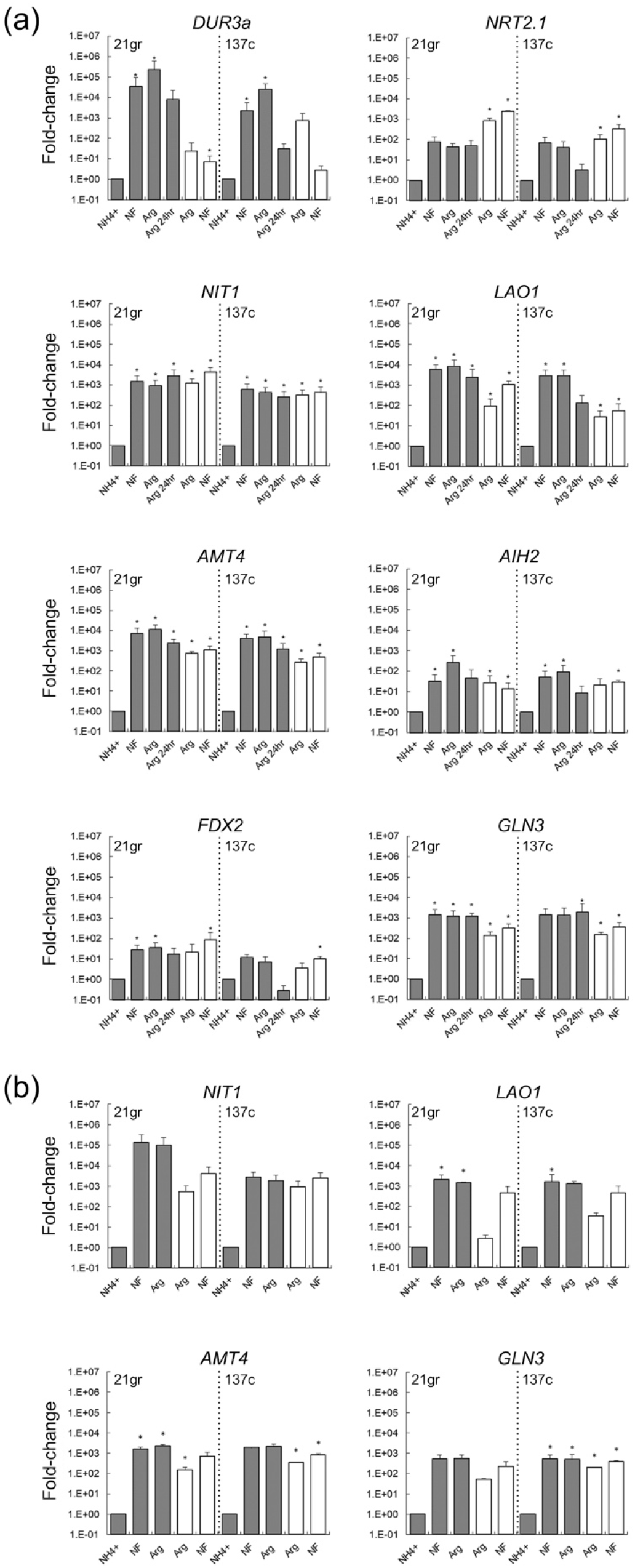
Arginine-grown cells constitutively upregulate N starvation marker genes. Expression of target transcripts relative to NH4+ condition. Ammonium (closed bars) or arginine (open bars) cultures of 21gr and 137c strains grown in mixotrophic condition (a) or in phototrophic condition (b) were harvested at mid-logarithmic phase and incubated in ammonium (NH4+), N-free (NF), arginine (Arg) medium for two hours or arginine medium for twenty-four hours (Arg-24h, only for mixotrophic cultures). Data represent the mean ± the standard deviation from three biological replicates. Asterisks indicate significance of the difference between the Arg or NF and the NH_4_+ data points using a Student’s t-test with P < 0.05.

To minimize acclimation responses to the change in N source, we examined arginine cultures pre-acclimated to the arginine medium for at least two growth cycles from logarithmic to stationary phase. The logarithmic culture of arginine-acclimated cells showed significant 27-to >1,200-fold upregulation of *AMT4, GLN3, LAO1, NIT1*, and *NRT2.1* and 3.5-to >500-fold upregulation of *DUR3A, FDX2, AIH2*, compared to ammonium cultures (Fig. 2a, Arg of arginine culture). When these arginine-acclimated cells were transferred to N-free medium, they exhibited only a modest 1.33-to 11-fold upregulation in all marker genes but *DUR3A*, whose expression was reduced close to the level in ammonium culture (Fig. 2a, N-free of arginine culture). Together, these results suggest that the same gene regulatory mechanism triggered by N starvation becomes activated during arginine growth regardless of genetic difference between 21gr and 137c (ex. *NIT1* and *NIT2*).

N starvation responses are generally considered as a stress response. We therefore asked whether arginine culture invokes additional cellular stress that may in turn activate N starvation responses. To assess physiological stress of arginine cultures, we analyzed chlorophyll *a* fluorescence parameters that are known to fluctuate upon cellular stress (Baker, 2008). Maximum quantum yield of PSII (F_v_ / F_m_) and photosynthetic efficiency (ΦPSII at 210 μmol photons m^−2^ s^−1^) in arginine cultures were not significantly different from those in ammonium cultures (Table 2b). The chlorophyll *a/b* ratio as an indicator of antenna size was found marginally higher in arginine cultures, significantly for 137c but not for 21gr. Arginine cultures showed a strong reduction in the cellular chlorophyll content, down 66% - 34% of the levels found in ammonium cultures and visibly evident as pale green logarithmic cultures (Table 2b; Fig. 3a). These results suggest that chlorophyll reduction may be due to an activated N starvation signal and not due to severe stress which would be evidenced as a drop in the F_v_ / F_m_ below 0.7 (Baker, 2008).

**Fig. 3.**
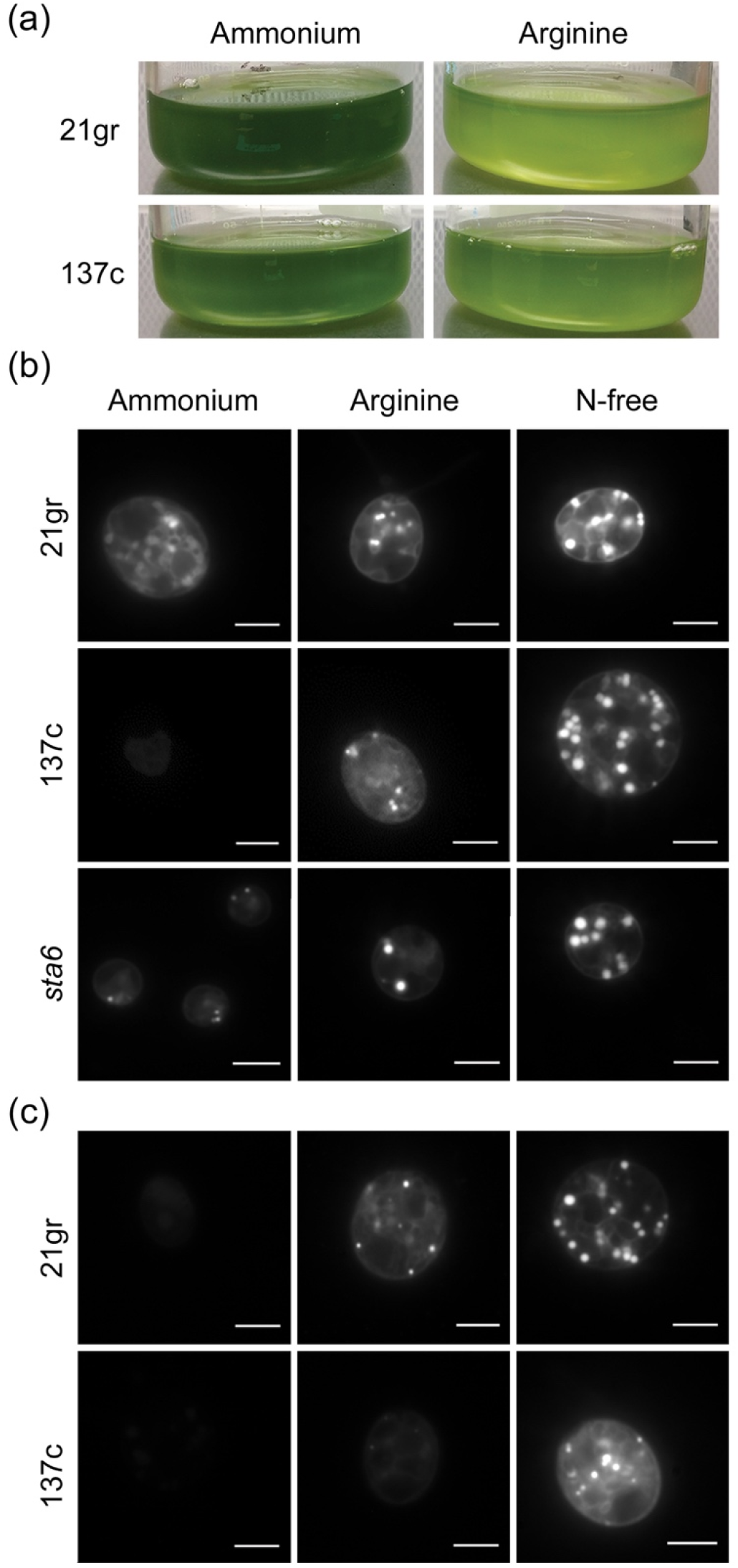
Arginine-grown cells exhibit N-starvation phenotypes. (a) Pale green logarithmic culture indicates reduced cellular chlorophyll content. Photographs of ammonium- or arginine-grown 21gr and 137c cultures were taken at their early logarithmic phase after adjusted to 2 x 10^6^ cells per mL. (b-c) Nile red staining reveals increased neutral lipid biosynthesis in cells grown with arginine. Micrographs of Nile red stained mid-logarithmic phase cells from ammonium or arginine cultures of 21gr, 137c, and *sta6* grown either mixotrophically (b) or phototrophically (c) alongside cells starved of N for 3 days captured using a 100x objective. The starchless mutant, *sta6* was used as a high lipid accumulating control line. Scale bar represents 5 μm.

### 3.2. Arginine-grown cells increase the flux of organic carbons towards storage

Lipid accumulation in response to N starvation is well documented (Hu et al., 2008; Wang et al., 2009); however, it is not known whether N starvation stimulates lipid body biogenesis directly or indirectly due to cell cycle arrest or generic stress responses. Lipid droplets were present in logarithmic arginine cultures with more pronounced accumulation in 137c (Fig. 3b). The relative size and number of lipid droplets found in the arginine cultures were less than those observed in cells starved of nitrogen for three days. Strain specific differences in lipid droplet formation could be due to differences in carbon flux towards starch biosynthesis that also increases during N starvation (Siaut et al., 2011).

To quantitatively assess the carbon flux into storage molecules without diversion into starch biosynthesis, we examined the *starchless* mutant, *sta6*, that shunts all storage carbons to lipids without apparent changes in TAG biosynthesis pathways (Ball et al., 1991; Fan et al., 2011; Li et al., 2010). The large number and size of lipid droplets in arginine-grown *sta6* cells attests to an increased carbon flux toward lipid storage molecules (Fig. 3b, *sta6).* In arginine cultures, the total fatty acid (FA) content per cell was 18% and 60% higher than the level in the ammonium-grown cells for 21gr and *sta6*, respectively (Table 4a). The TAG content per cell also increased 27% and 317% from the level in ammonium culture (Table 4b). For *sta6*, TAGs accounted for 28% of the cellular FA content in the arginine-grown cells, more than 2-fold greater than the ammonium-grown cells (13%). The arginine-grown wild-type 21gr cells, however, showed no significant enrichment of TAG, attesting diversified carbon flux in wild-type cells. This result suggests increased FA biosynthesis occurs in arginine cultures, as reported for the N-starved *C. reinhardtii* cells (Moellering and Benning, 2010).

**Table 4.**
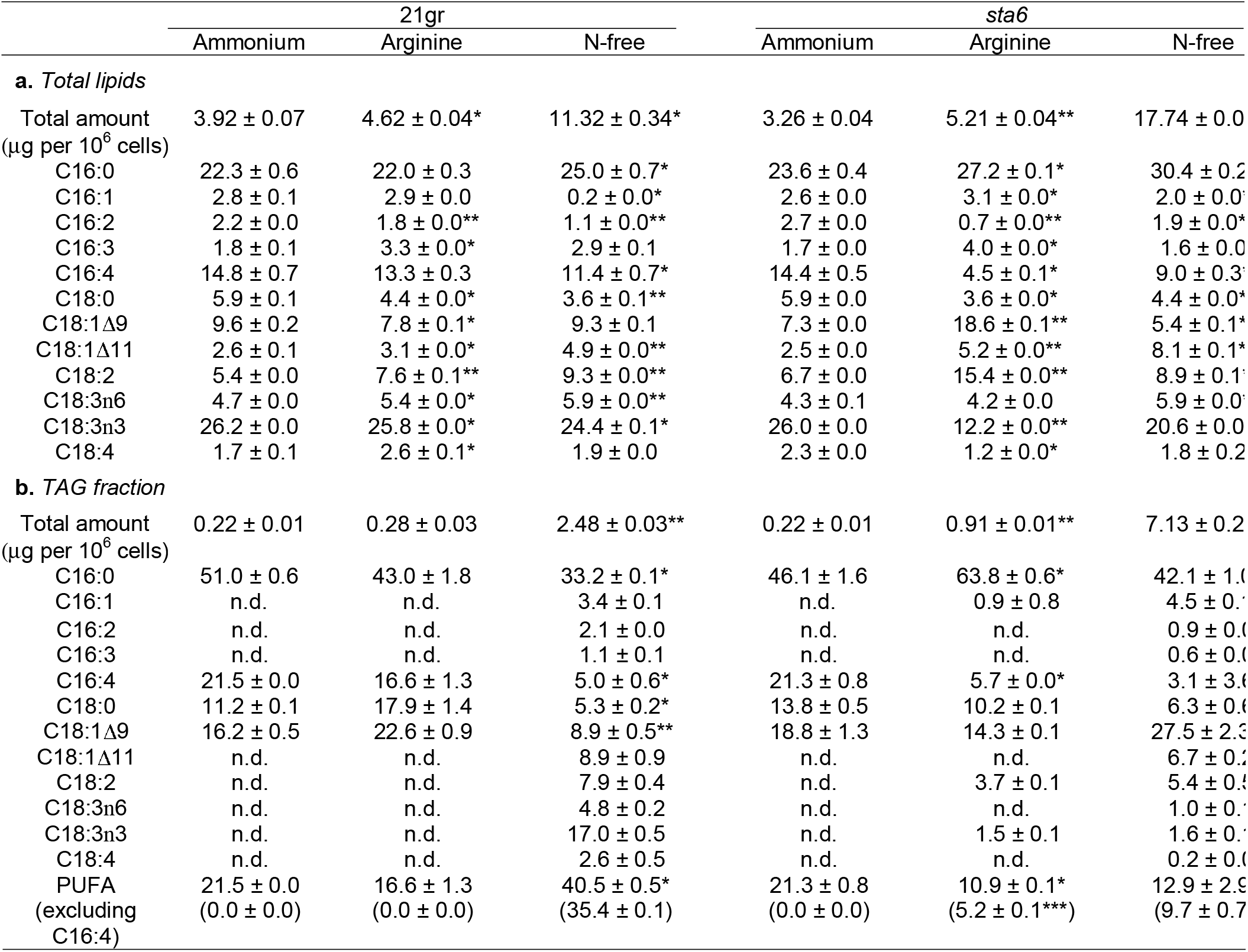
FA profile suggests *de novo* synthesis as a primary route for the increased FAs in arginine culture. Relative quantity (%) of FA profile in total lipid extracts (a) or in TAG fractions (b) was analyzed using gas chromatography of mixotrophic cultures of 21gr and *sta6* grown to mid-logarithmic phase using either ammonium or arginine, alongside ammonium-grown cells starved of N for three days (N-free). FAME quantification of total lipids and TAG fractions is provided in the row (Total amount), expressed in μg per 10^6^ cells, above the FA profile data. Values are means ± standard deviation (%, n >=2). Significant difference from ammonium data is indicated with an asterisk (P <0.05) or double asterisks (P <0.01) and was calculated by a Student’s t-test. Δ indicating the position of the double bond counting from the carboxyl end and n indicating the position from the terminal methyl carbon. n.d., not detected; PUFA, polyunsaturated fatty acids; ***, significant difference from N-free data (P <0.01)

Increased incorporation of FA into TAG can have two explanations: *de novo* synthesis and recycling of membrane lipids (Boyle et al., 2012; Fan et al., 2011; Liu et al., 2013; Moellering and Benning, 2010; Siaut et al., 2011). To investigate the sources of increased FAs, we analyzed the relative quantity of FA profiles distinguishing saturated and monounsaturated FAs, representing newly synthesized fatty acids, and polyunsaturated FAs (PUFAs) such as C16:4, C18:3, and C18:4, representing typical forms of membrane lipids (Ohlrogge and Browse, 1995). FA profiles of total lipids within arginine cultures of 21gr and *sta6* were significantly different from those of ammonium cultures, showing an increase in C16:0 and C18:1Δ11 (Δ indicating the position of the double bond counting from the carboxyl end) and a decrease in C16:4, which was comparable to the FA profiles of N-starved cultures except an increase of C18:1Δ11 instead of C18:1Δ9 (Table 4a). The FA profiles of the TAG fraction showed that the increased cellular TAG content of the *sta6* arginine culture was contributed primarily by a 574% increase in C16:0 and a >300% increase in C18:0 and C18:1Δ9. PUFA content of the TAG fraction showed a significant reduction in the *sta6* arginine culture from that of N-starved *sta6* cells, and the reduction was 46% when excluding C16:4, the major FA of photosynthetic membrane (Giroud et al., 1988) (Table 4b). The low quantity of PUFAs indicates a minor flux of membrane lipid recycling into TAG biosynthesis in arginine cultures. Our results suggest that the increased TAG in arginine culture depends on *de novo* FA synthesis.

### 3.3. N starvation responses are induced in phototrophic arginine cultures

N starvation signaling presumably operates in all trophic states, whereas some N starvation responses may depend on the trophic state: for example, decreases in chlorophyll content during N starvation in *C. reinhardtii* were observed only in the mixotrophic state but not in the phototrophic state (Terauchi et al., 2010). To test whether the N starvation responses of arginine cultures depend on the trophic state, we also examined phototrophic arginine cultures. *AMT4, GLN3*, and *NIT1* were up-regulated by 53- to >500-fold, comparable to the mixotrophic arginine cultures, in both 21gr and 137c (Fig. 2b). *LAO1* was 33-fold upregulated in 137c but not in 21gr. Lipid droplets were present in phototrophic arginine cultures of 21gr strain, whereas they were few or absent in those of 137c strain (Fig. 3c). The growth rates of phototrophic arginine cultures were 27-37% lower than those of phototrophic ammonium cultures (Fig. 1b, Table 3b). However, the stationary phase culture density was 3.2 to 6.9-fold higher in arginine cultures (Table 3b). We found no significant difference between ammonium and arginine phototrophic cultures in chlorophyll *a* fluorescence parameters (Table 2c). The chlorophyll content of phototrophic arginine culture was also comparable to that of phototrophic ammonium culture, suggesting that the ~50% decreased chlorophyll content in mixotrophic arginine culture is due to the activated N starvation responses and trophic state rather than any arginine-dependent effects. Collectively, phototrophic arginine culture also showed N starvation responses without any apparent defect in photosynthesis and growth. This observation is important for biofuel development since mixotrophic cultures are more prone to bacterial/fungal contamination than phototrophic cultures. The three to seven-fold increase in the final cell density recorded in the phototrophic arginine cultures is of great interest towards the aim of improved biomass yield (Cheirsilp and Torpee, 2012). Whether higher cell density is associated to constitutive N starvation response of cells or to any other arginine-specific effects warrants a further investigation.

### 3.4. Arginine-induced lipid accumulation occurs in the diatom Phaeodactylum

Arginase is a major enzyme for arginine catabolism in plants and algae; however, the *C. reinhardtii* genome lacks genes encoding this enzyme (Vallon and Spalding, 2009; Winter et al., 2015). To test whether arginine culture can elicit N starvation responses in photosynthetic organisms with arginase, we examined a marine diatom, *P. tricornutum*, that can grow with arginine as the sole N source and expresses arginase (Allen et al., 2011; Fig. 4a). As in *C. reinhardtii*, pronounced lipid droplets were found in *P. tricornutum* when growing solely with arginine for seven days (Fig. 4b). A fluorescence-based semi-quantitative assay demonstrated a two-to seven-fold higher neutral lipid content during a seven-day growth period (Fig. 4c). Similar lipid accumulation suggests that the putative mechanism triggered by arginine cultures may be conserved across diverse photosynthetic organisms.

**Fig. 4.**
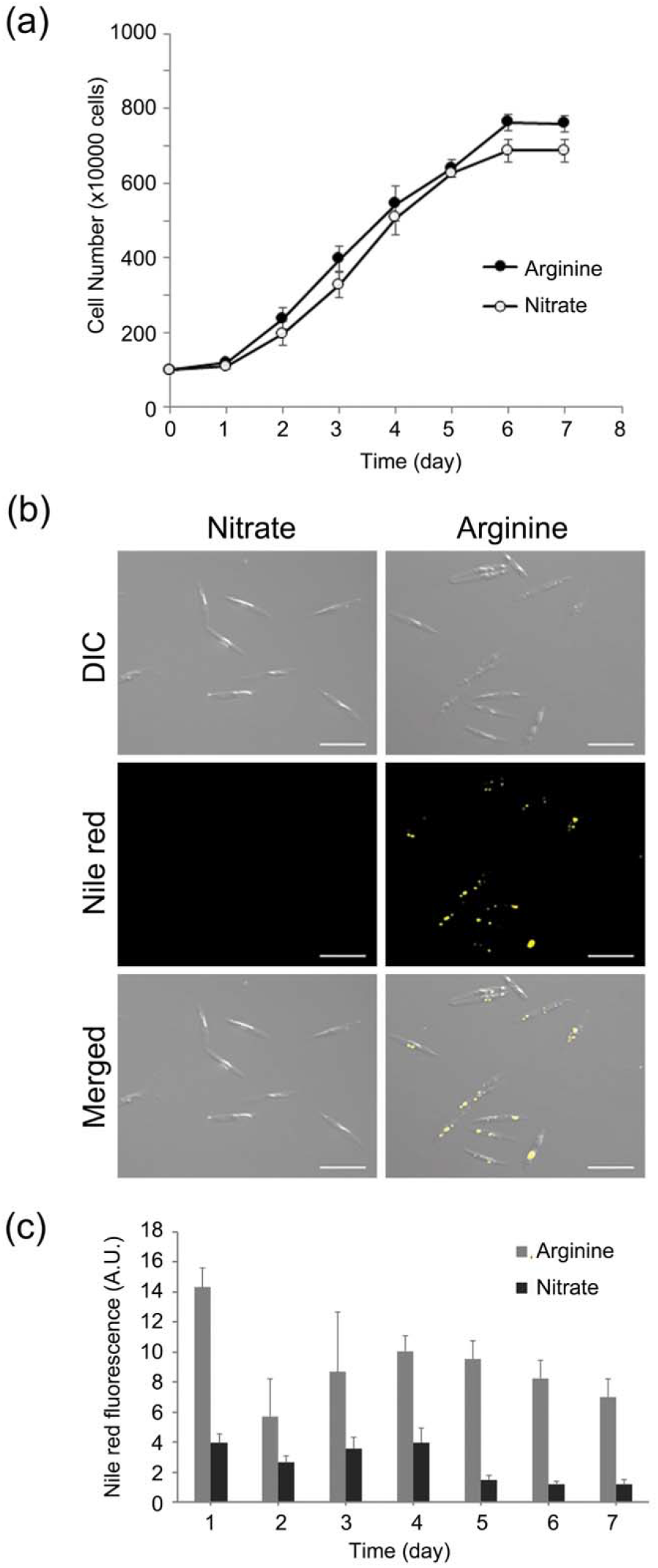
Arginine culture of *P. tricornutum* increases neutral lipid content during robust growth. (a) Culture proliferation measured by cell counting with a hemocytometer for cultures grown with arginine (closed circles) or nitrate as the sole N source (open circles). Data represent the mean and standard deviation from triplicate sampling in two independent experiments (n=6). (b) Micrographs of Nile red stained cells from cultures grown for seven days with nitrate or arginine as the sole N source grown. Bar = 20 μm. (c) Semi-quantitative determination of total neutral lipid accumulation per 10^6^ cells using a fluorescence-based assay comparing overall Nile red fluorescence between nitrate and arginine cultures. Error bars are standard deviation from triplicate sampling in two independent experiments (n=6). Significant differences between arginine and nitrate cultures were found in all the time points when calculated by a Student’s t-test (P < 0.05). A.U., arbitrary unit.

## 4. Conclusion

Arginine cultures of *C. reinhardtii* in mixotrophic and phototrophic conditions activate a suite of N catabolite genes together with N starvation-induced responses such as gametogenesis and TAG accumulation while supporting growth to a comparable or higher maximum density than ammonium cultures. We documented a similar induction of TAG accumulation in arginine cultures of a diatom, *P. tricornutum.* Hence, arginine feeding serves to alleviate canonical N catabolite repression by ammonium or nitrate and provides a fruitful approach to manipulate N starvation responses without growth arrest in microalgae concomitant with the goal of developing practical solutions to improve biodiesel productivity and quality.

## Acknowledgments

This work was supported by Discovery Grant 418471-12 from the Natural Sciences and Engineering Research Council (NSERC) (to J.-H.L.), by the Korea CCS R&D Center (KCRC), Korean Ministry of Science, grant nos. 2016M1A8A1925345 (to J.-H.L.), 2014M1A8A1049278 (to S.J.S.) and 2014M1A8A1049273 (to E.J.).

## Declarations

There are no potential financial or other interests that could be perceived to influence the outcomes of the research. No conflicts, informed consent, human or animal rights applicable. All authors confirmed the manuscript authorship and agreed to submit it for peer review.

## Author contributions

Munz, J. designed and performed the experiments, analyzed the data, and wrote the manuscript. Xiong, Y. performed the experiments and analysis of the photosynthesis. Kim, J.Y.H. and Sung, Y.J. performed the experiments and analysis of lipid extraction and FAME quantitation. Seo, S. performed the experiments and analysis of *P. tricornotum.* Hong, R.H. and Lee, J. contributed to the RNA extraction and the RT-qPCR. Kariyawasam T. contributed to the microscopic observation and analysis. Shelly N. performed the photobioreactor experiments. Sim, S.J. contributed to the reviewing and editing of the manuscript in addition to supervising FAME analysis part of the research. Jin, E.S. contributed to the reviewing and editing of the manuscript in addition to supervising *P. tricornotum* part of the research. Lee, J.-H. designed the experiments, analyzed the data, and wrote the manuscript.

## Abbreviations

FA: fatty acid
FAME: fatty acid methyl ester
N: nitrogen
PUFA: polyunsaturated fatty acid
TAG: triacylglycerol
TAP: Tris-acetate-phosphate medium

